# Using targeted therapy to promote a pro-inflammatory tumour microenvironment and anti-tumour immune response in high grade serous ovarian cancer

**DOI:** 10.1101/2025.08.10.669554

**Authors:** Zhen Zeng, Anastasia Gandini, Rituparna Bhatt, Martina Proctor, Nicole-Lisa Li-Ann Goh, Shivam Vora, Thomas P. Walsh, Sherry Y. Wu, Kaltin Ferguson, Jermaine I. Coward, Snehlata Kumari, Nikolas K. Haass, James W. Wells, Janet Hardy, Lewis Perrin, Yaowu He, John D. Hooper, Gwo-Yaw Ho, Jazmina L. Gonzalez Cruz, Brian Gabrielli

## Abstract

**BACKGROUND:** High-grade serous ovarian cancer (HGSOC) is characterized by elevated replication stress and an immunosuppressive microenvironment. A synergistic combination of checkpoint kinase 1 inhibitor (CHK1i) with low-dose hydroxyurea (LDHU) promotes a unique ATR-independent moderate replication stress response with potent anti-tumour effects. The ability of this approach to reprogram the tumour immune microenvironment (TIME) to overcome the immunosuppression and promote an anti-tumour immune response in HGSOC is the focus of this study.

**METHODS:** We investigated the therapeutic potential of CHK1i+LDHU in established HGSOC cell cultures, fresh tumour cell explants from HGSOC patient ascites, and syngeneic mouse models, assessing tumour cell killing, immunogenic cell death, pro-inflammatory cytokine/chemokine expression, and anti-tumour immune responses.

**RESULTS:** CHK1i+LDHU effectively killed ovarian cancer cells regardless of chemotherapy resistance, *BRCA2* mutation and homologous recombination repair status *in vitro*. *In vivo*, treatment significantly reduced tumour burden and ascites accumulation. CHK1i+LDHU enhanced expression of pro-inflammatory cytokines/chemokines and triggered immunogenic cell death in tumour. In syngeneic models, treatment promoted CD8^+^ cytotoxic T cell-dependent anti-tumour responses and reduced immunosuppressive signalling within the TIME.

**CONCLUSIONS:** CHK1i+LDHU is a promising therapy for chemotherapy-resistant HGSOC, combining direct cytotoxic effects with reprogramming the TIME to reduce immunosuppression and activate a CD8^+^ T cell-dependent anti-tumour response.

## Background

High grade serous ovarian cancer (HGSOC) is the most lethal gynaecologic malignancy, with a 5-year survival rate of 35-40% ^1^. The current standard of care involves platinum-based chemotherapy, often combined with taxanes. While initially effective, over 70% of patients experience relapse within 2 years ^2^. Chemotherapy is associated with high-grade toxicities that severely impact patient quality of life and, importantly, contributes to immunosuppression ^3^. Relapse-free survival is extended in patients with homologous recombination repair defective (HRD) tumours (< 50% of patients) treated with poly-ADP ribose polymerase inhibitors (PARPi) ^2,4^. However, this benefit is limited in non-BRCA mutant ovarian cancer, and emerging data indicate increased toxicity associated with PARPi use in the relapse setting ^5^. Immunotherapy response rates are <20% ^6^, underscoring the profound immunosuppressive barriers in HGSOC ^7^. This low response rate is particularly surprising given that a high proportion of HGSOC cases exhibit tumour-infiltrating lymphocytes, and CD8^+^ T cell levels are prognostic for survival ^6^. This indicates the tumour immune microenvironment (TIME) is highly immunosuppressive, severely limiting the efficacy of immunotherapies ^8^. Elevated levels of regulatory T cells (Tregs), M2 type macrophages and myeloid derived suppressor cells (MDSCs) are commonly observed in HGSOC, contributing to this immunosuppressive TIME ^9^. Consequently, once patients develop resistance to chemotherapy, treatment options are severely limited. There is an urgent need for targeted therapies that not only selectively treat chemo-resistant HCSOC but also modulate the TIME to overcome the immunosuppression and promote robust anti-tumour immune responses.

Checkpoint kinase 1 inhibitors (CHK1i) effectively target cells with elevated replication stress ^10–12^, a common feature of ovarian cancer ^13^ and has been identified as an early feature of the disease ^14^. CHK1i have been shown to be effective in PARP inhibitor resistant HGSOC preclinical models ^12^. The CHK1i Prexasertib achieved a 30% disease control rate but and overall response rate of 10% in a phase II trial in platinum refractory HGSOC patients ^15,16^. Prexesertib combination with the PARPi Olaparib demonstrated effectiveness preclinically and in a phase I trial in PARPi resistant patients but was associated with significant haematological toxicity ^17–19^. Replication stress inducers such as gemcitabine combine synergistically with CHK1i ^20–22^. Clinical trials of these combinations across various cancer types (>12 trials), many in the recurrent, relapsed, or refractory setting ^23^, demonstrated anti-tumour efficacy but also revealed frequent severe haematological toxicities ^24,25^. These toxicities arise from CHK1i chemo-sensitising all tissue to the replication stress inducers which promote high-level replication stress in both tumour and normal tissue ^26^.

Subclinical doses of hydroxyurea (low-dose HU; LDHU) that have little effect on tumour growth either *in vitro* or *in vivo* ^10^ synergise strongly with CHK1i *in vitro* and *in vivo* in other cancer types ^10,27^. Unlike clinically used replication stress inducers such as gemcitabine that trigger an ATR/CHK1-dependent S phase checkpoint arrest ^28^, LDHU triggers a unique CHK1-dependent reorganisation of replication origin firing ^29^. LDHU selectively sensitises tumours with elevated endogenous replication to CHK1i at doses ∼20% of its reported Cmax ^10,30^. CHK1i are unlikely to be clinically viable as single agents, while most of the combinations trialled to date have excessive toxicity. This makes the combination of CHK1i+LDHU an attractive option owing to its efficacy, low normal tissue toxicity *in vivo* and its ability to maintain robust adaptive immune response ^10,31^.

In this study, we have investigated whether low dose hydroxyurea elicits the same synergy with CHK1i SRA737 in HGSOC as reported in other cancer types. Using a cross section of different molecular subtypes and treatment resistant HGSOC models, we assessed whether the broad tumour-killing efficacy observed elsewhere extends to HGSOC. Importantly, given the highly immunosuppressive TIME in HGSOC – which underlies its poor response to current immunotherapies – we also examined whether this combination can not only spare immune function but actively trigger anti-tumour immune responses. Specifically, in HGSOC, we investigated whether tumour cell death induced by CHK1i+LDHU is immunogenic, whether it remodels the cytokine and chemokine milieu to reprogram the TIME, and whether it overcomes key immunosuppressive barriers. We further identified the critical immune cell population mediating the treatment related anti-tumour effect. We demonstrate that CHK1i+LDHU efficiently kills a broad range of genotypes and chemo-resistant ovarian cancer cells. *In vivo*, this combination significantly reduced tumour burden and ascites accumulation by inducing immunogenic tumour cell death and activating a CD8^+^ T cell-dependent immune response, accompanied by a favourable reprogramming of the TIME. Together, these finding highlight CHK1i+LDHU as a promising therapeutic strategy for HGSOC, with the potential to overcome its profound immunosuppressive barriers and stimulate effective anti-tumour immunity.

## Methods

### Cell lines

Luciferase-tagged mouse ovarian cancer cell lines ID8-p53^WT^ and ID8-p53^−/−^ ^32,33^, human ovarian cancer cell lines OVCA420 (RRID:CVCL_3935), PEO1 (RRID:CVCL_2686), PEO4 (RRID:CVCL_2690), FUOV-1 (RRID:CVCL_2047), OVCAR3 (RRID:CVCL_0465), OVCAR8 (RRID:CVCL_1629) and Kuramochi (RRID:CVCL_1345) were derived as described in Supplementary Table S1. A panel of four cell lines, PEO1, OVCAR8, FUOV-1 and Kuramochi representing primary and metastatic deposits, BRCA wild type and mutant, chemo naive and resistant were used in all experiments. Other cell lines were added to this panel to examine the diversity expected in a normal patient population. Cells were cultured in high glucose DMEM (Gibco) with 10% heat treated fetal bovine serum (FBS; Bovogen), 1 mM Sodium pyruvate (Gibco), GlutaMAX (Gibco), 20 mM HEPES (Sigma-Aldrich) and antibiotic-antimycotic. Cell cultures were maintained in a Binder low oxygen incubator at 37°C, 5% CO_2_ and 2% O_2_ and were tested mycoplasma free.

### Patient ascites-derived tumour cells

Tumour cells were isolated from ovarian cancer patient ascites (as described in Supplementary Table S2) through centrifugation at 340 g for 10 minutes, followed by red blood cell lysis with ACK buffer (10 mM Tris-Cl, 5 mM MgCl_2_, 10 mM NaCl). The isolated cells were resuspended and cultured for 24 hours in complete DMEM media. This initial culture allowed most of the fibroblast population to adhere. Subsequently, all media was transferred into a new flask and cultured for an additional 48 hours to allow tumour cell adherence. The media was then changed to a supplemented mixture consisting of 33% Medium 199 (Gibco), 33% DMEM/F12 (Gibco), 33% RPMI (Gibco) with 10% heat-treated fetal bovine serum (FBS; Bovogen), 1% Insulin-Transferrin-Selenium (Gibco), 25 ng/ml Cholera Toxin (Sigma-Aldrich), 0.5 µg/ml Hydrocortisone (Sigma-Aldrich), 10 ng/ml Epidermal Growth Factor (Sigma-Aldrich), 10 ng/ml Fibroblast Growth Factor (Sigma-Aldrich), and antibiotic-antimycotic. After reaching confluency, differential trypsinization was performed to remove any remaining fibroblasts by adding 0.25% Trypsin-EDTA (Gibco) for 40 seconds and removing the supernatant containing the fibroblasts ^34^. Cell cultures were maintained in a Binder low oxygen incubator at 37°C, 5% CO_2_, and 2% O_2_.

### Dose-response assay

All cell lines were seeded in 96-well plates and treated with increasing concentrations of CHK1i (SRA737, Sierra Oncology) in combination with 0.2 mM HU (Sigma). Cells were assessed for viability after treatment for 3 days using resazurin (Sigma).

### Time-lapse cell viability killing assay

Indicated cells (500-3000 in 100ul media per well) were seeded into 96-well plates and treated with or without 1 µM CHK1i (SRA737), ATR inhibitor 2 μM VE-821 (SelleckChem) and 0.2 (low dose) or 2 mM (high dose) hydroxyurea. Sytox Green (125 nM; Molecular Probes) to mark dead cells was also added and cells were imaged every four hours at 10x magnification for up to 6 days using an IncuCyte S3 live cell imaging system (RRID:SCR_023147). Analysis was performed using IncuCyte software (RRID:SCR_025367) to identify total number of cells, and percentage of cells positive for Sytox Green staining for each frame. Data was presented as total cell counts for proliferation, percent cell death from the number of Sytox stained cells/ total cell number for each time point, and percent viable cells.

### Immunoblotting

Ovarian cancer cell lines were treated with or without 1 µM CHK1i (SRA737) and 0.2 mM hydroxyurea for 24 hours. Cell pellets were lysed and analysed by immunoblotting, using antibodies against RPA2 (Cat#52448, RRID:AB_2750889), γH2AX (Ser139/Tyr142) (Cat#5438, RRID:AB_10707494) (Cell Signalling Technology) and Alpha-Tubulin (Cat#600-401-880, RRID:AB_2137000) (Rockland). Antibodies were diluted 1:1000. Proteins were visualized using chemiluminescence (Fusion SL Viber Lourmat).

### RT-qPCR

Murine (ID8-p53^WT^ and ID8-p53^−/−^) and human (PEO1, PEO4, OVCAR8, OVCA420, FUOV-1 and Kuramochi) ovarian cancer cell lines were treated with CHK1i combination (1µM CHK1i and 0.2mM hydroxyurea) for 24 hours. Total RNA was extracted using TRIzol (Invitrogen) per the manufacturer’s protocol. RNA concentration and purity were assessed using a NanoDrop One/OneC. cDNA synthesis was performed with 2 µg of RNA using the High-Capacity cDNA Reverse Transcription Kit (Applied Biosystems: 4368814), followed by a 1:10 dilution with nuclease-free water. Quantitative PCR was conducted using PowerUp SYBR Green Master Mix (Applied Biosystems) on a ViiA 7 Real-Time PCR System or QuantStudio 7 Flex Real-Time PCR System. Custom and predesigned KiCqStart Primers (MERK) were employed for qPCR. The genes analysed are listed in Table S3. Fold change in gene expression was calculated using 2^–(ΔCt), where ΔCt represents the difference between the average Ct of the target gene and the reference gene, actin (mouse) or YWHAZ (human). <u>Cytokine Bead Array</u> LEGENDplex HU Essential Immune Response Panel (BioLegend, San Diego, CA, USA) was used according to the manufacturer’s instructions to assess the cytokines present in cell supernatants after 48 h CHK1i+LDHU treatment. Samples were performed in duplicate and analysed on a CytoFLEX S Flow Cytometer, RRID:SCR_019627 (Beckman Coulter, Lane Cove, Australia). Acquired data were analysed using provided LEGENDplex Data Analysis Software.

### ICD Marker

Human ovarian cancer cell lines were cultured for 24 h in the presence of DMSO or CHK1i +LDHU, then harvested and stained for surface expression with anti-calreticulin-APC, anti-HSP90-PE or relevant isotype controls (diluted in 1:100). The delta mean-fluorescence-intensity (ΔMFI) values were calculated by taking the mean values on live cells. Stained cells were analysed using an LSR-Fortessa X20 Flow Cytometer (BD BioSciences) with FACSDiva software, RRID:SCR_001456 (Becton Dickinson, Sparks, MD, USA). Acquired data were analysed using FlowJo software, RRID:SCR_008520 (TreeStar Inc., Ashland, OR, USA).

### Immune Profiling

Tumours in omentum were harvested, minced and treated with DNASe1 and collagenase IV for 30 min, then pushed through a 40 μm cell strainer to generate a single-cell suspension. Cells were blocked using Fc block then stained with Live/dead Aqua (Thermo Fisher Scientific, Waltham, MA, USA) and a panel of conjugated antibodies in 1:100 dilution for immune cell profiling (CD45.2-PE dazzle, CD3-FITC, TCRβ-PE, CD8α-BV605, CD4-BUV395, NK1.1-PE-Cy7, CD11b-BV421, F4/80-BV711, Gr-1-PercpCy5.5, Ly6G-AF700, MHCII-APC-Cy7, CD19-BV785). FoxP3 (FoxP3-AF647) was detected using the FoxP3 Staining Kit (eBioscience) as per the manufacturer’s instructions. In some experiments, the lymphoid and myeloid markers were separated, PD-L1BV711 and CD86-APC added to the myeloid panel and CD25-BV421 added to the lymphoid panel. Flow-Count Fluorospheres (Beckman Coulter, Miami, FL, USA) were used for total cell counts. Single Ab-stained compensation beads were used to set the gating. Stained cells were analysed using an CYTEK Aurora Spectral Flow Cytometer (Cytek Biosciences) with SpectroFlo software, RRID:SCR_025494. Acquired data were analysed using FlowJo software, RRID:SCR_008520.

### Mouse tumour assays

Experiments were performed with approval from The University of Queensland Animal Ethics Committee (2021/AE000249). Luciferase-labelled ID8-p53^WT^ and ID8-p53^−/−^ cells (1.5 × 10^6^) in 200 μl of Hank’s balanced salt solution were intraperitoneally (i.p.) injected into C57BL6/J mice (RRID:IMSR_JAX:000664). Tumour establishment was confirmed using an IVIS Lumina X5 imaging system, with luciferin bioluminescence images acquired and analysed by in vivo imaging software. Following tumour establishment, mice were treated with CHK1i (SRA737)+LDHU as described previously ^31^. To examine the influence of CD8^+^ T cells on treatment responses, anti-CD8β (Lyt 3.2, BioXcell) or appropriate isotype control injections were initiated 3 days prior to CHK1i+LDHU treatments and then repeated weekly. At the experimental endpoint, ascites fluid was collected, and tumours in omentum and other organ sites were dissected in a double-blinded manner.

### Statistical analysis

All statistical analyses were performed using GraphPad Prism 9 (RRID:SCR_002798). Bar graphs display mean values and standard deviation.

## Results

We assessed the efficacy of the CHK1i+LDHU combination across a panel of established mouse and human ovarian cancer cell lines and in tumour cells recently derived from chemo-resistant HGSOC patient ascites. The panel is representative of chemo-sensitivity, *BRCA* mutation and homologous recombination deficiency (HRD) status found in HGSOC patients (details in Table S1 and S2). Dose-response assays using the clinically tested CHK1 inhibitor (SRA737) combined with 0.2 mM HU revealed all tested cell lines displayed similar sensitivity to the CHK1i+LDHU combination, with IC_50_ values below 1 μM for CHK1i (Figure 1A). This is significantly lower than the reported 6 μM highest concentration (Cmax) in patients ^30^. Based on these findings, a combination of 1 μM CHK1i and 0.2 mM HU was selected. The synergy of this combination was assessed in three cell lines. As single agents, CHK1i and LDHU treatment had little or modest effects on cell proliferation and cell killing, whereas the combination blocked proliferation and promoted high levels of cell killing in the tested cancer cell lines (Figure 1B). This clearly demonstrated the individual components had little anti-proliferative effect, and the synergy of the combination. The degree of cell killing achieved by CHK1i+LDHU was in most cases greater than CHK1i combination with 2mM HU (HDHU) (Supplementary Figure S1). Cell killing was calculated as percentage of dead cells of the total cell number in each frame and underrepresents level of the cell killing relative to the either starting cell number or control cell numbers at the same time point. By calculating the percentage of viable cells in the final frame in the control and treated samples relative to the first frame of the time lapse sequence, treated samples had final cell viability >40% of the initial cell number across a larger panel of human and murine OvCa lines (Figure 1C, Supplementary Figure S2). The sensitivity did not appear to be determined by the genotype or chemo-sensitivity as the most sensitive cell lines (those with lowest viable cell count with treatment) OVCAR3 and OVCAR8 were both chemo-resistant, and BRCA deleted and wild type, respectively (Table S1). LDHU alone produced a modest reduction in proliferation (Figure 1B), differentiating it from the ATR-dependent checkpoint response triggered by agents such as HDHU ^35^. ATR inhibitor had no effect when combined with LDHU and had modest effect on cell killing when combined with HDHU (Figure 1D), similar to that observed with CHK1i+HDHU (Supplementary Figure S1). Tumour cells cultured from patient ascites samples (GO579, GO618, GO623) were also sensitive to the combination (Figure 2), demonstrating similar sensitivity to CHK1i+LDHU in recent patient derived cells. Together, these data demonstrate that CHK1i+LDHU effectively induces cell death across a range of ovarian cancer models, irrespective of genetic background, HRD status, or chemotherapy resistance.

**Figure 1:**
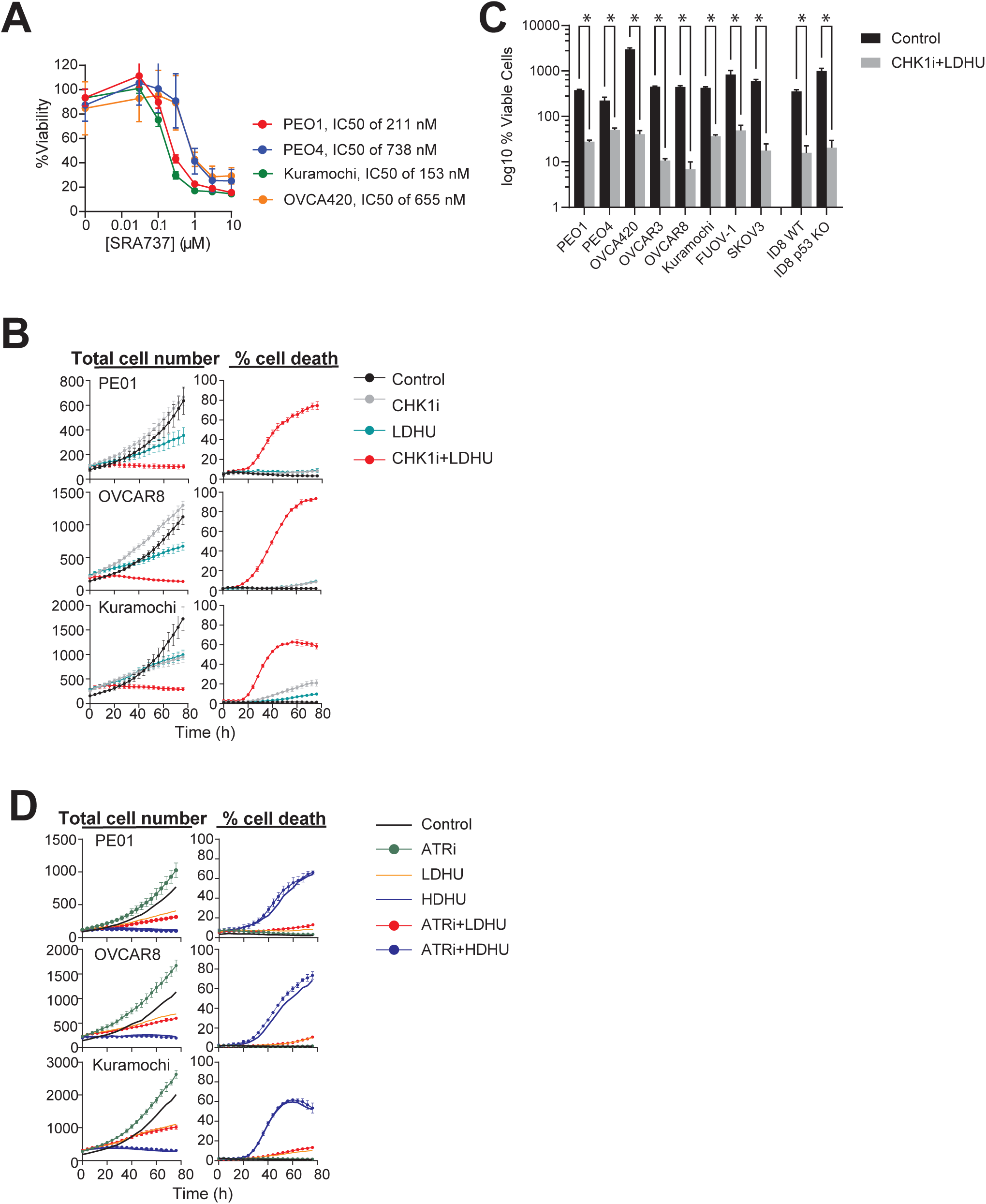
Ovarian cancer cell lines are sensitised to CHK1i combination. **A)** Dose–response curves of the indicated ovarian cancer cell lines treated with CHK1i (SRA737) in the presence of a constant 0.2 mM hydroxyurea (HU). Cell viability was assessed using a resazurin assay. Data represent the mean ± SD of 4–5 replicates and are representative of duplicate experiments. **B–D)** The indicated HGSOC cell lines (see Tables S1), were treated under the following conditions: Control (vehicle), CHK1i (1 µM SRA737), LDHU (low-dose HU, 0.2 mM), HDHU (high-dose HU, 2 mM), ATRi (2 µM VE-821), CHK1i+LDHU, ATRi+LDHU, and ATRi+HDHU. Cells were monitored using the IncuCyte system for up to five days. Total cell numbers and % cell death we calculated as outlined in the Methods. % viable cells was calculated from number of viable (Sytox negative) cells in final time point of the Incucyte experiments/cell number in the initial time point. Data represent the mean ± SD of 6–30 replicates from 3 independent experiments. * p<0.0005 using multiple t-test.

**Figure 2:**
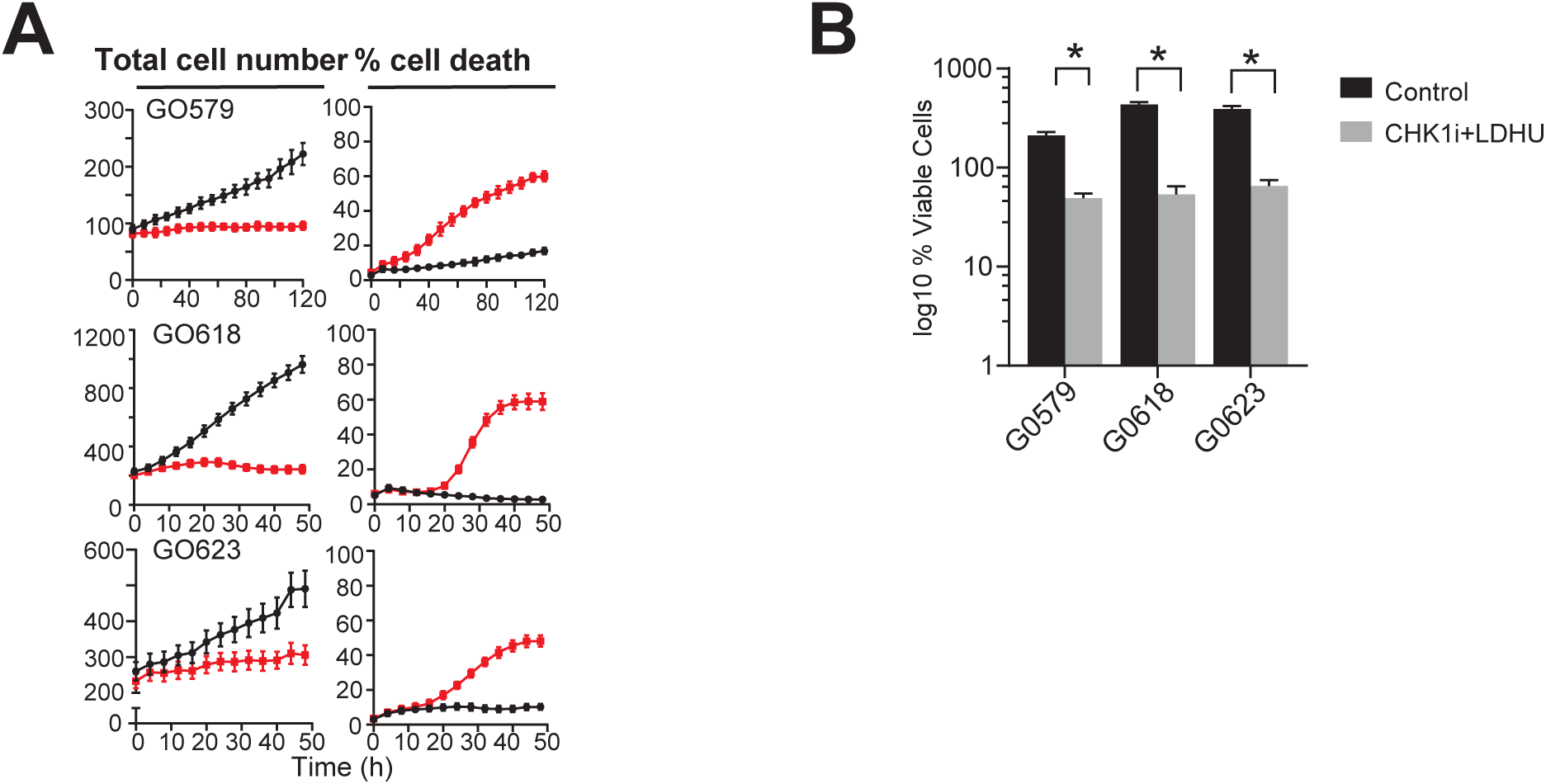
Primary ovarian cancer cells are sensitised to CHK1i combination. **A**) The indicated HGSOC primary cells derived from patient ascites samples (see Tables S2), were treated with Control (vehicle) or CHK1i (1 µM SRA737) plus LDHU (low-dose HU, 0.2 mM), Cells were monitored using the IncuCyte system for up to five days. **B)** % viable cells was calculated from number of viable (Sytox negative) cells in final time point of the Incucyte experiments/cell number in the initial time point. Data represent the mean ± SD of 6–30 replicates from 3 independent experiments. * p<0.001 using multiple t-test.

CHK1i+LDHU induced replication stress in ovarian cancer cells, as evidenced by the slower migrating RPA2 band that is indicative of RPA2S4/8 phosphorylation ^36^, and increased DNA damage, indicated by elevated γH2AX levels in the tested cell lines (Supplementary Figure S3), consistent with previous findings ^10^. DNA damage is known to increase pro-inflammatory cytokine and chemokine expression ^37^. CHK1i+LDHU treatment significantly increased the expression of pro-inflammatory signals *CCL5*, *CXCL8*, *CXCL10, IL-6,* and *TNF* at RNA levels across all human ovarian cell lines (Figure 3A). In murine ovarian cancer lines, *CCL5*, *CXCL10*, and murine equivalents of *CXCL8* (*CXCL1* and *CXCL2*) were similarly upregulated (Figure 3B). Expression of anti-inflammatory *TGFb* and *VEGFA* were also modestly increased with treatment in the human cell lines (Figure 3A). The changes in levels of the secreted proteins (Figure 3C) generally mirrored the RNA expression patterns (Figure 3A), with pro-inflammatory signals (CCL5, CXCL8, CXCL10, IL-6 and TNFα) increasing with treatment, but the free form of TGFβ was unchanged. Overall, CHK1i+LDHU promoted a pro-inflammatory microenvironment, upregulating expression of CCL2, CCL5, CXCL10 and TNFα while leaving the anti-inflammatory cytokine TGFβ largely unaffected.

**Figure 3:**
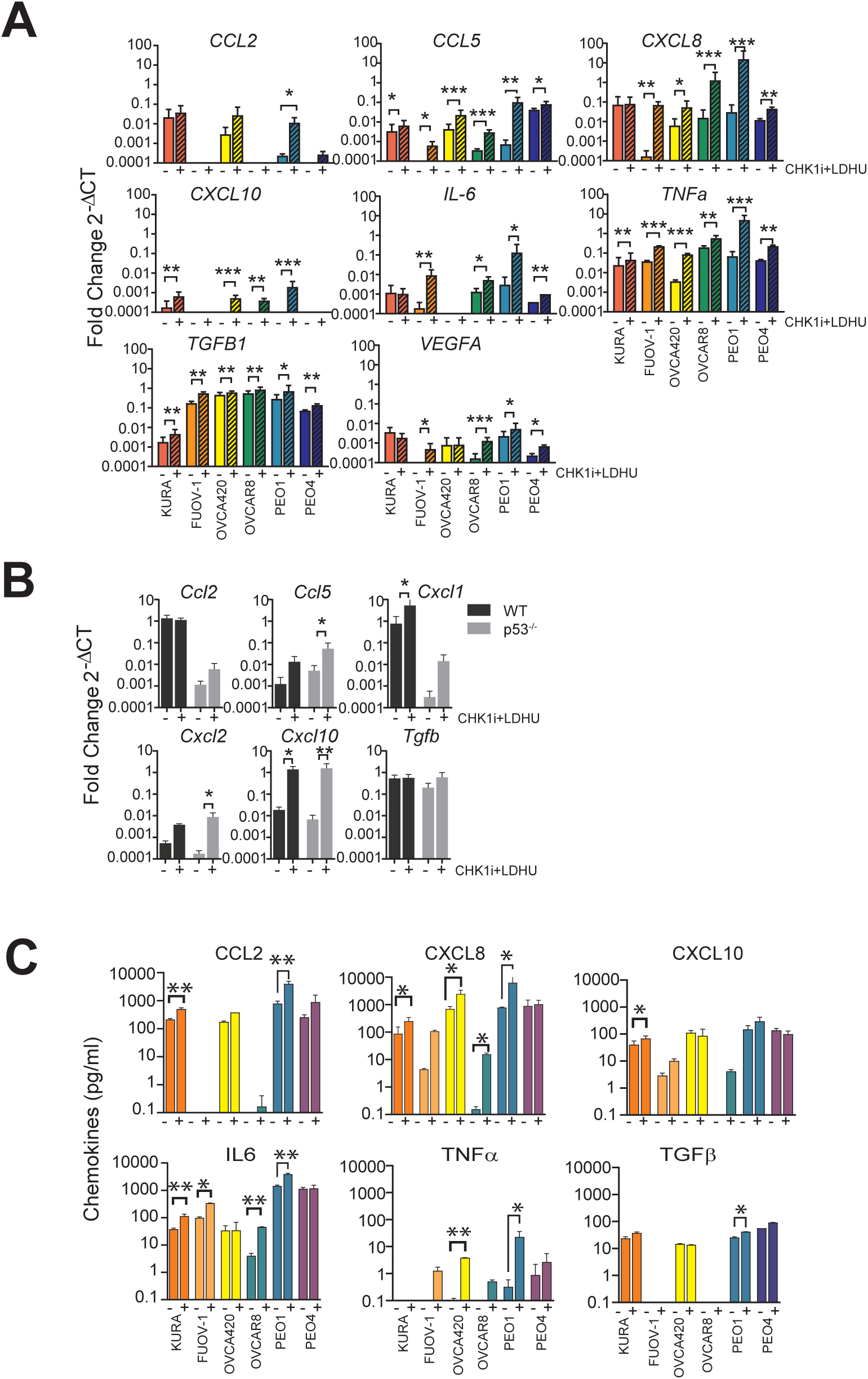
CHK1i combination treatment induces proinflammatory responses in ovarian cancer cells. The indicated ovarian cancer cell lines were treated with or without 1 µM CHK1i (SRA737) + 0.2 mM HU for: **A, B)** 24 hours, followed by cell harvesting and measurement of the mRNA for indicated cytokines/chemokines using qRT-PCR, and **C)** 48 hours, followed by harvesting supernatants for cytokine level measurement. The levels of the qRT-PCR products are shown relative to control. The qRT-PCRF data are representative of 2-4 independent experiments with at least three replicates. Chemokine levels were measured in 2 individual experiments with 2 replicates. P values calculated by t-test for each chemokine. * p<0.05, ** p<0.01, *** p<0.001.

In addition to its direct cytotoxicity and induction of pro-inflammatory signals, we investigated whether CHK1i+LDHU treatment could further modulate the TIME by triggering immunogenic cell death (ICD). The cell surface expression of damage-associated molecular patterns (DAMPs) calreticulin and HSP90 were assessed in a representative panel of cell lines. There was significantly increased surface expression of calreticulin in OVCAR8 and Kuramochi, and increased surface HSP90 in PEO1 and Kuramochi, although increased total expression of both proteins was observed in all cell lines with treatment (Figure 4A). To determine the immunogenic potential of CHK1i+LDHU *in vivo*, luciferase-labelled mouse ID8-p53^−/−^ cells treated *in vitro* with CHK1i+LDHU for 24 h were used to inoculate immunocompetent mice. Freeze-thawed tumour cells were used as a negative control for ICD response. After 9 days, all mice were challenged with live, untreated ID8-p53^−/−^ tumour cells injected into the opposite flank and tumour growth was monitored (Figure 4B). Tumour burden, assessed by bioluminescence imaging, was similar in the PBS control and freeze-thawed cell inoculated mice, whereas mice inoculated with CHK1i+LDHU-treated ID8-p53^−/−^ cells were significantly protected (Figure 4C).

**Figure 4:**
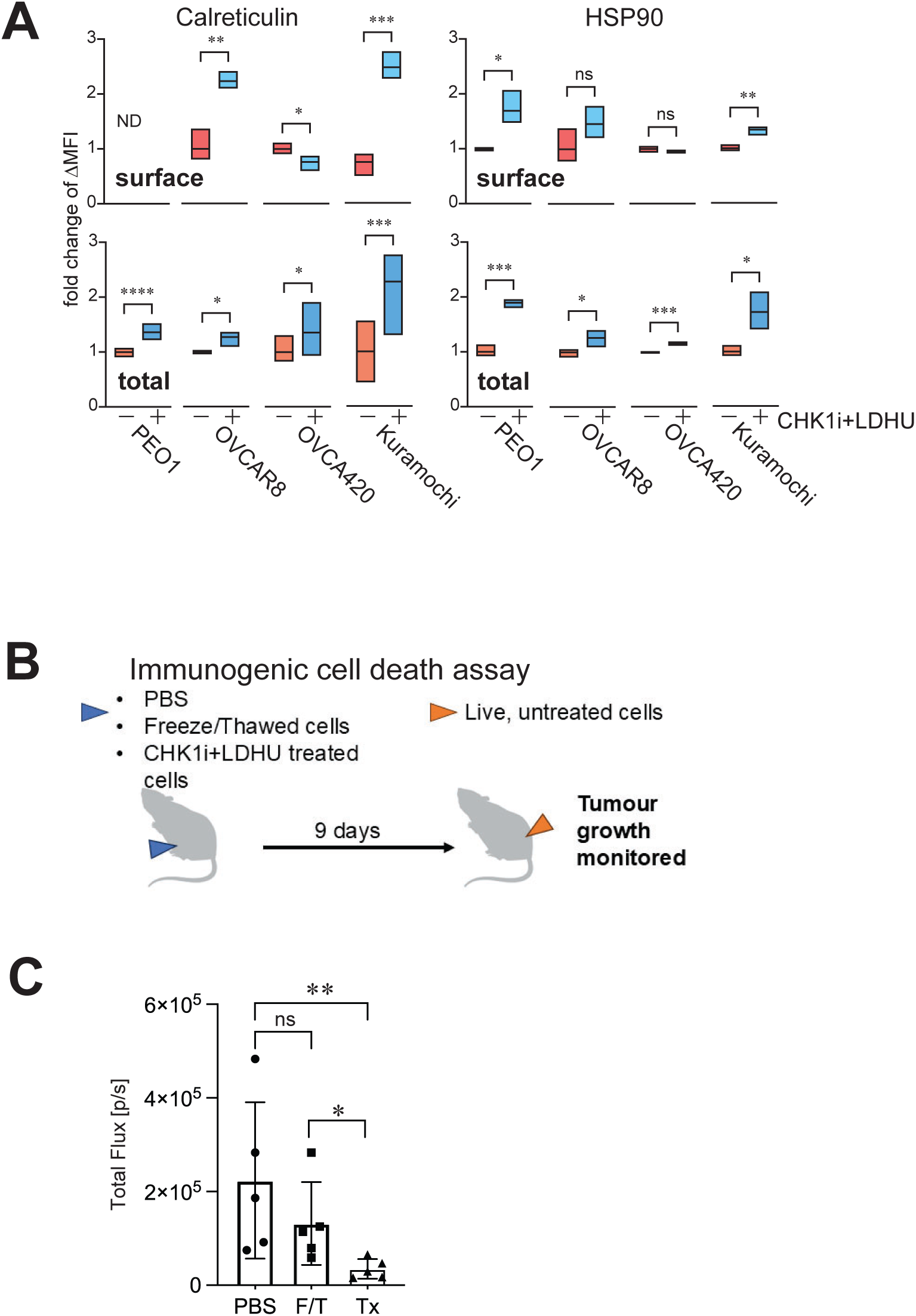
CHK1i combination treatment promotes ICD in ovarian cancer models. Human ovarian cancer cell lines were treated with or without 1 μM CHK1i (SRA737) + 0.2 mM HU and harvested at 24 h for analysis of calreticulin and HSP90 on live cells. **A)** Fold change in surface (top) or total (bottom) expression levels of calreticulin and HSP90. Data are representative of two independent experiments. ND, none detected. **B)** C57BL/6J mice were immunised with ID8-p53^−/−^ cells treated in vitro 1 µM SRA737 + 0.2 mM HU (Tx) for 24 h. Freeze-thaw killed (F/T) ID8-p53^−/−^ cells were used as a negative control for ICD. At 9 days after immunisation mice were rechallenged with live ID8-p53^−/−^ cells into the opposite flank and tumour burden was monitored by IVIS in vivo. N= 5 mice each treatment. Statistical analysis using unpaired t-test * p<0.05, ** <0.01, ***<0.001, ****<0.0001.

Based on these finding, we further investigated the *in vivo* therapeutic efficacy of CHK1i+LDHU. Two different murine ovarian cancer models, luciferase-labelled ID8-p53^WT^ and ID8-p53^−/−^ tumours were assessed using syngeneic C57BL/6J mice. Three weeks of treatment with 3 days/week CHK1i+LDHU (Figure 5A) effectively blocked ascites accumulation and tumour growth in both models (Figure 5B,C) but had little effect on mouse body weights (Figure 5D). The slightly lower body weights of the treated mice accounted for the reduced ascites and tumour weight. To assess whether the pro-inflammatory cytokine and chemokine expression triggered by the treatment altered the TIME, immune profiling was performed using a panel of flow cytometry markers for myeloid and lymphoid cell types. In ID8-p53^WT^ tumours, immunosuppressive MDSCs and Tregs were significantly decreased, accompanied by an increase in macrophage abundance (Figure 6A). An extended panel of markers was used to investigate NK cells in the ID8-p53^−/−^ model. Additionally, the immune cell composition of the ascites in this model was assessed. Changes similar to the ID8-p53^WT^ model myeloid compartment were observed in the ID8-p53^−/−^ tumours (Figure 6B). The decreased MDSC and Treg in the tumour was mirrored by the changes in these cell types in the ascites. The proportion of CD8^+^ T cells in the CD45^+^ leukocytes was greater in this model than the ID8-p53^WT^ tumours, but there were only minor changes in the abundance of CD8^+^ T cells with CHK1i+LDHU treatment in either model. The reduction in the CD4^+^ T cells abundance reflected the reduction in the Treg levels (Figure 6A, B). Notably, a Ly-6C expressing CD8^+^ T cell subset (detected by the Gr-1 antibody and more cytotoxic than normal effector T cells ^38^) was significantly increased with treatment, and Ly-6C expression was 3-fold higher in this subset compared to non-treated tumours (Figure 6C).

**Figure 5:**
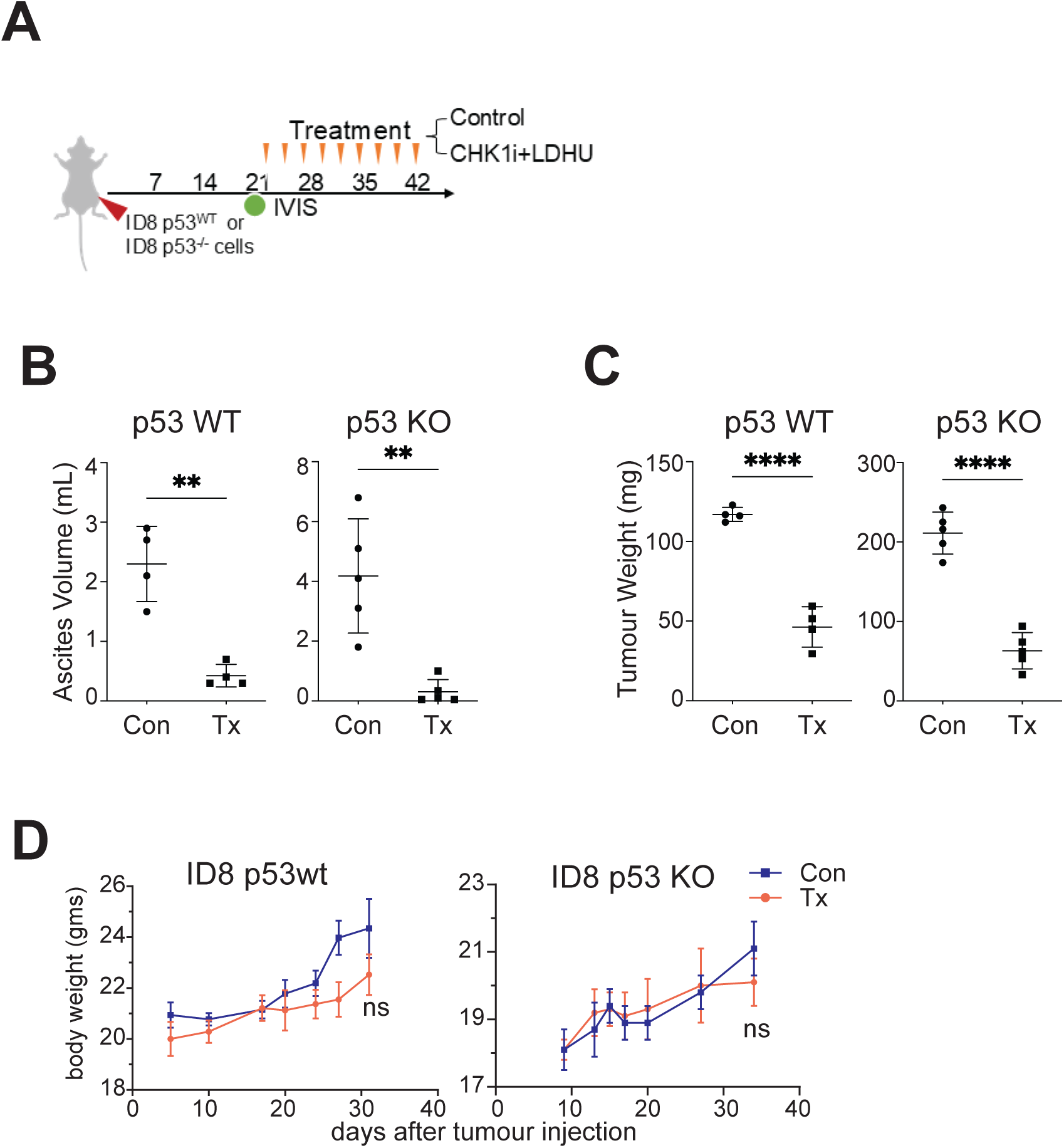
CHK1i combination treatment effectively controls ascites accumulation and tumour growth of ID8-p53^WT^ and ID8-p53^−/−^ models. Syngeneic mouse ovarian cancer models ID8-p53^WT^ and ID8-p53^−/−^ were established in immunocompetent mice (n=5 per treatment group). Once tumour engraftment and dissemination throughout the abdominal cavity were confirmed by IVIS imaging, mice were treated with vehicle control or CHK1i combination therapy, as previously described ^31^. **A)** Volume of ascites, and **B)** tumour burden at endpoint in mice treated (Tx) or untreated (Con) with three weekly cycles of CHK1i combination therapy. **C)** Body weight measurements following tumour cell injection over the course of the study. Statistical analysis using unpaired t-test ** p<0.01, ****<0.0001.

**Figure 6:**
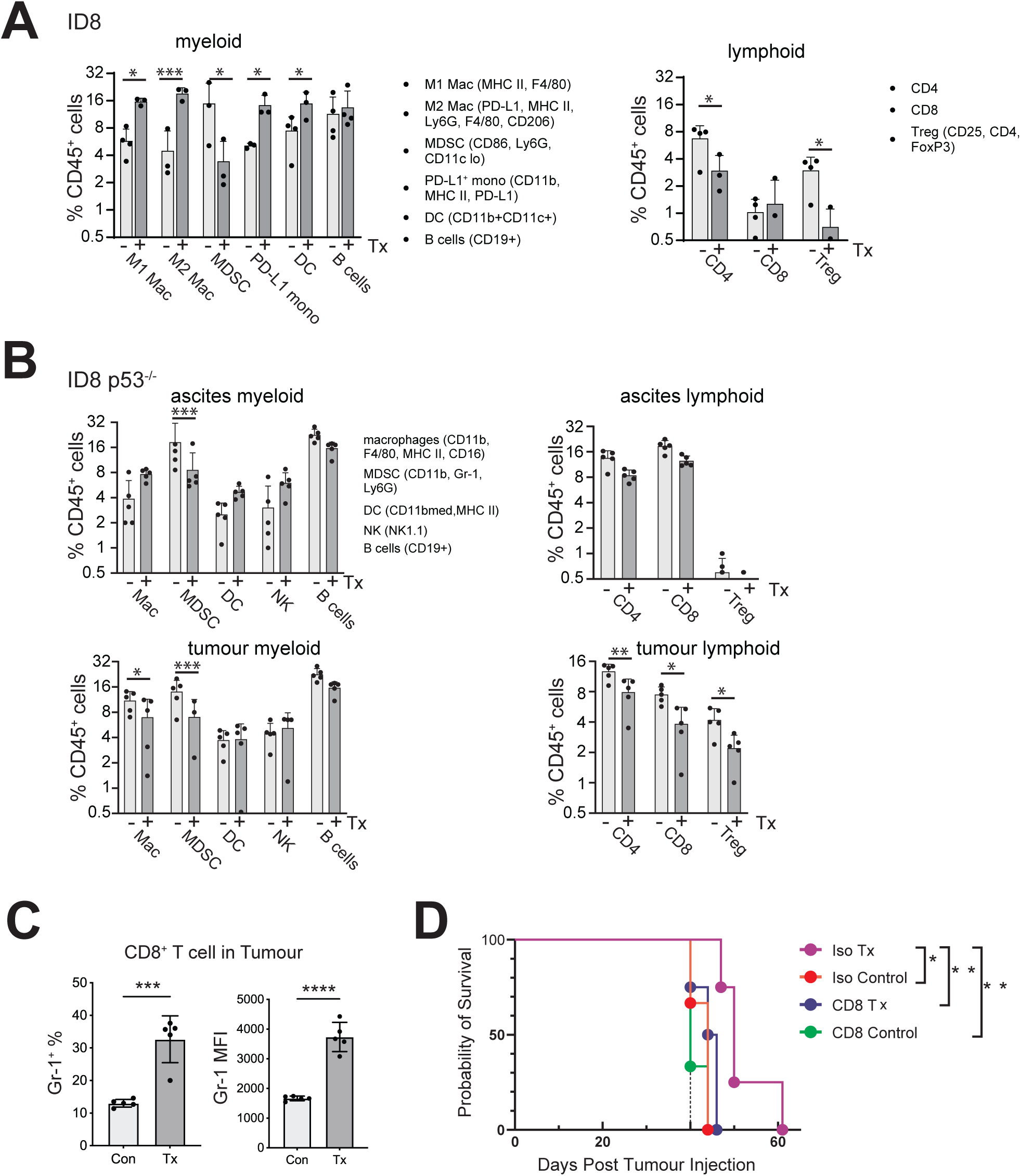
CHK1i combination induced anti-tumour immune response is dependent on CD8^+^ T cells. ID8 and ID8 p53^−/−^ tumours were established in immunocompetent mice. Once tumour transplantation has been confirmed by IVIS, treatment as in Figure 5 was commenced (day 21). **A, B)** Ascites and tumour samples (omentum part) collected from Figure 5 were processed and stained with 13 immune cell markers for cell phenotype analysis using Cytek Aurora Flow Cytometry System. The relative abundance of the indicated major immune cell types in either the myeloid (MDSCs, macrophages, DCs, NK cells, B cells), or lymphoid cell compartments (CD4^+^ and CD8^+^ T cells, Tregs) was assessed using the markers shown. **C)** Gr-1expression on CD8^+^ T cells (left: Percentage of Gr-1^+^ cells gated from CD8^+^ T cells; right: MFI of Gr-1 on CD8^+^ T cells) in ID8-p53^−/−^ tumours. **D)** In ID8-p53^−/−^tumour bearing mice, antibody injections, either isotype control or anti-CD8β were initiated on the same day CHK1i+LDHU treatment (day 21) and then repeated on day 28 and day 35 along with the normal CHK1i+LDHU treatment protocol. Kaplan-Meier graph showing the survival of mice treated with: Iso = Isotype mAb, CD8 = anti-CD8 mAb, Control = Placebo control, Tx = CHK1i+LDHU. Statistical analysis using unpaired t-test * p<0.05, ** <0.01, ***<0.001, ****<0.0001. For survival, used log rank test.

To determine whether cytotoxic CD8^+^ T cells are involved in the CHK1i combination induced anti-tumour responses, CD8^+^ T cells were specifically deplete in the ID8-p53^−/−^ model using anti-mouse CD8β (Lyt 3.2, Clone 53-5.8). The 53-5.8 antibody has been shown to achieve complete depletion of CD8^+^ T cells *in vivo* without affecting CD8^+^ CD11c^+^ dendritic cells which primarily express the CD8αα homodimer ^39^. Depletion of CD8^+^ T cells using 53-5.8 MAB (Supplementary Figure S4) abolished the ability of CHK1i+LDHU to control the tumour growth, demonstrating a key role for CD8^+^ T cells in the combination’s mechanism of action (Figure 6D).

## Discussion

Current chemotherapy for HGSOC is initially effective but will inevitably lead to resistance and comes with very significant toxicity to the patients ^40^. Here we demonstrate that a synergistic combination of subclinical doses of CHK1i and low dose HU is effective in killing a high proportion of HGSOC cell lines and new patient derived models *in vitro*, irrespective of genotype, HRD status or chemo-sensitivity. The broad effectiveness of the CHK1i+LDHU combination has also been observed in melanoma and NSCLC models ^10^. A combination of CHK1i with low dose gemcitabine has also been shown to be effective in SCLC where it also triggers a CD8^+^ T cell dependent immune response ^41^. However, even low doses of gemcitabine can trigger the ATR/CHK1-dependent S phase checkpoint arrest rather than the unique ATR-independent, CHK1-dependent replication origin reorganisation that is triggered by LDHU ^10,26^. Notably, unlike the higher concentrations of HU or gemcitabine that are effective as single agents and trigger an S phase arrest, the low dose HU used here trigger a CHK1i-dependent response without causing growth arrest either *in vitro* or *in vivo* ^10,27,29^. Of note, normal tissue, especially the proliferating cell in the colonic crypts and immune cells that are responsible for the many toxic side effects of chemotherapy are not adversely affected by CHK1i+LDHU *in vitro* or *in vivo* ^10,31^. The broad effectiveness of CHK1i+LDHU is likely to be due to replication stress inherent in the development of HGSOC ^14^.

The synergistic CHK1i+LDHU combination can promote a pro-inflammatory environment that is the result of both pro-inflammatory cytokine/chemokine expression and DAMPs released as a consequence of tumour cell ICD ^42^. Targeted therapies such as PARP inhibitors can also trigger ICD and pro-inflammatory responses ^42,43^, but are associated with high rates of high-grade haematological toxicity including reduction in lymphocytes ^44^, suggesting that the immunomodulation by PARPi is negated by a sub-optimal immune response. This may account for the lack of efficacy of combination of PARPi with immunotherapy ^44^. The relative lack of deleterious effect of CHK1i+LDHU on either T cell proliferation *in vitro* or on a T cell dependent immune response *in vivo* ^31^ provides strong evidence that this combination has minimal negative impact on the immune function, making an anti-tumour immune response a viable outcome.

Overcoming the immunosuppressive state common in ovarian cancer requires an anti-tumour treatment capable of targeting multiple components. Beyond its direct tumour cell–killing effects, CHK1i+LDHU induced expression of pro-inflammatory chemokines and cytokines such as CCL2, CCL5, CXCL10 and TNF-α. This overall shift towards a pro-inflammatory environment may have played a pivotal role in modulating immune cell recruitment and activity. CXCL10 has been shown to recruit and restimulate T cells, including effector CD8^+^ T cells ^45^. Similarly, CCL2 and CCL5 have been implicated in the recruitment of antigen-presenting cells including dendritic cells (DCs) to inflammatory sites, further supporting their role in enhancing antigen presentation and T cell activation ^46^. High CCL2 levels promote a pro-inflammatory Th1/Th17 phenotype, while CCR2 deficiency shifts the immune balance toward Tregs ^47^. High CCL5 expression promotes the recruitment of CD8^+^ T cells, NK cells, and activated macrophages ^48^, and CCL5 has been shown to recruit CCR5^+^ dendritic cells to enhance CD8^+^ T cell priming and sustain anti-tumour responses ^49^. CHK1i+LDHU significantly reprogrammed the TIME by reducing immunosuppressive MDSCs and Tregs - critical barriers to effective immune surveillance in HGSOC ^50^, and by activation of CD8^+^ T cell depletion was surprising given. We demonstrated that much of the treatment induced anti-tumour response was dependent on CD8^+^ T cells. The marked loss of tumour control following CD8^+^ T cells were depleted was surprising as the given the combination’s efficacy in killing tumour cells *in vitro*. Similarly, treatment response was lost in syngeneic melanomas grown RAG1^-/-^ mice which lack adaptive immunity ^31^, despite being highly effective in immunocompromised Nude mice ^10^. These findings suggest that the anti-tumour effect induced by CHK1i+LDHU is not solely due to direct tumour cell killing but also relies on CD8⁺ T cells, providing strong evidence that this treatment harnesses the immune system to combat the tumour – an effect that is particularly important in HGSOC, where most patients do not respond to immunotherapy.

In this study, we demonstrated that CHK1i+LDHU induces a robust anti-tumour immune response, modulating both innate and adaptive immune responses. Current immunotherapies used as single agents are effective in tumours that have already triggered an immune response, the so called “hot” tumours, but are ineffective in “cold” tumours where no prior immune response exists ^51^. The ability of CHK1i+LDHU to induce ICD and enhance expression of pro-inflammatory cytokines/chemokines reshapes the TIME, reducing immunosuppressive elements such as MDSCs and Tregs, and importantly, establishes a potent CD8^+^ T cell-mediated anti-tumour response regardless of its initial status. A limitation of this study is that all *in vivo* experiments were performed in single mouse strain, C57-BL/6J, which essentially mimics a single patient in terms of the genetic diversity of the immune response.

In conclusion, synergistic CHK1i+LDHU combination therapy represents a promising strategy for treating HGSOC by directly killing tumour cells and importantly, reprogramming the immune microenvironment to promote an effective anti-tumour immune response *in vivo*. By including primary tumour cells derived from patient ascites alongside a diverse panel of established human ovarian cancer cell lines, this study provides strong evidence that the efficacy of CHK1i + LDHU extends to clinically relevant, heterogeneous tumour contexts. The high sensitivity in diverse HGSOC models, even in chemo-resistant tumours suggests that the CHK1i combination could be an effective option for patients who have exhausted standard treatment options.

## Acknowledgments

We thank patients and staff who contributed to the Mater Research Biobank. We acknowledge the support of Mater Misericordiae Hospital and Gynaecological Oncology Unit in the collection of the Clinical Subject Data and Clinical Subject Materials. We also acknowledge the support of our consumer representatives Jacinta Frawley, Wanda Lawson and Lynelle Armitage. ID8-p53WT, ID8-p53^−/−^ and Kuramochi lines were kindly provided by Professor Roby (Kansas University Medical Center), Professor McNeish (Imperial College London), and Professor Konecny (University of California Los Angeles), respectively.

## Author Contributions

Conceptualization, ZZ, BG.; methodology, ZZ, RB, SYW, YH, JLGC; validation, ZZ, AG, RB, NLLAG, NJAR.; formal analysis, ZZ, SYW, AG, NLLAG, RB, JLGC, BG.; investigation, ZZ, AG, RB, MP, NLLAG, SV,TPW, SYW, JLGC; resources, SYW, KF, SK, NKH, JH, LP, YH, JDH, GYH.; data curation, ZZ, NLLAG,.MP; writing—original draft preparation, ZZ, AG, BG.; writing—review and editing, ZZ, AG, RB, MP,NLLAG,SYW, NKH, JWW, JDH, GYH, JLGC, BG.; visualization, ZZ, AG, NLLAG, BG.; supervision, ZZ, RB, SYW, JLGC, BG.; project administration, BG.; funding acquisition, JLC, JLGC, BG. All authors have read and agreed to the published version of the manuscript.

## Ethics approval and consent to participate

Human ethics was provide by the Mater Misericordiae Ltd Human Research Ethics Committee (MML HREC) for Project Id: 29596, approval PRGRPT/MML/29596. Informed consent was obtained from each patient for their ascites samples, and the study was performed in accordance with the Declaration of Helsinki. Animal ethics was obtained from The University of Queensland Molecular Biosciences Animal Ethics Committee – 2021/AE000249.

## Data Availability Statement

The original contributions presented in this study are included in the article/supplementary material. Further inquiries can be directed to the corresponding author(s).

