## Supplementary material for "Using targeted therapy to promote a pro-inflammatory tumour microenvironment and anti-tumour immune response in high grade serous ovarian cancer"

### Supplementary Information

#### Tables

Supplementary Table S1: Characteristic of the ovarian cancer cell lines used.

| Cell line | Disease | Tissue Derived from | Treatment history | BRCA1/2 status | Additional mutations |
|---|---|---|---|---|---|
| **FUOV-1** | High grade ovarian serous adenocarcinoma | Primary tumour | Chemo naïve | WT | TP53 mutated |
| **Kuramochi** | High grade ovarian serous adenocarcinoma | Ascites | N/A | BRCA2 mut | TP53 mutated |
| **OVCAR8** | High grade ovarian serous adenocarcinoma | Primary tumour | Progression on Carboplatin | WT | TP53, KRAS, ERBB2, CTNNB1 |
| **OVCAR3** | High grade ovarian serous adenocarcinoma | Ascites | The patient previously received cyclophosphamide, cisplatin, and doxorubicin treatment | BRCA2 deleted | TP53 mutated |
| **OVCA420** | Ovarian serous adenocarcinoma | Primary tumour | N/A | WT | TP53 mutated |
| **PEO1** | Ovarian cystadenocarcinoma | Ascites | The patient previously received cisplatin, 5-fluoruoracil and chlorambucil treatment and developed clinical resistance to chemotherapy. | BRCA2 mut | TP53 mutated |
| **PEO4** | Ovarian cystadenocarcinoma | Ascites | Another cell line derived from the same patient as PEO1 | BRCA2 revertant | TP53 mutated |
| **SKOV-3** | Ovarian serous cystadenocarcinoma | Ascites | N/A | WT | TP53 deleted |
| **ID8 p53 WT** | | Mouse primary tumour | | WT | None |
| **ID8 p53 KO** | | Mouse primary tumour | | WT | TP53 deleted |

Supplementary Table S2:

| Patient number | Disease | Treatment history | BRCA status | HRD status | Additional mutations |
|---|---|---|---|---|---|
| GO579 | HGSOC | Carboplatin and Paclitaxel | No pathogenic | Positive | TP53 |
| GO618 | HGSOC | Unknown | No pathogenic | Negative | TP53 |
| GO623 | HGSOC | Carboplatin and Paclitaxel | Unknown | Unknown | Unknown |

Supplementary Table S3: RT-qPCR primers

| Primer (Mouse) | Sequence (5'-3') | Primer (Human) | Sequence (5'-3') |
|---|---|---|---|
| **FM1 ACTIN** | CGAATCATGAGCATTGTAGAC | **FM1 CCL2** | AGACTAACCCAGAAACATCC |
| **BM1 ACTIN** | GTAATTCTTATCTCCAGCCAG | **BM1 CCL2** | ATTGATTGCATCTGGCTG |
| **FM1 CCL2** | CAAGATGATCCCAATGAGTAG | **FM1 CCL5** | ACTTGCCTCCCCATATTC |
| **BM1 CCL2** | TTGGTGACAAAAACTACAGC | **BM1 CCL5** | AAGAGTTGATGTACTCCCG |
| **FM1 CCL5** | AGGAGTATTTCTACACCAGC | **FM1 CXCL8** | GTTTTTGAAGAGGGCTGAG |
| **BM1 CCL5** | CAGGGTCAGAATCAAGAAAC | **BM1 CXCL8** | TTTGCTTGAAGTTTCACTGG |
| **FM1 CXCL1** | AAAGATGCTAAAAGGTGTCC | **FM1 CXCL10** | AAAGCAGTTAGCAAGGAAAG |
| **BM1 CXCL1** | GTATAGTGTTGTCAGAAGCC | **BM1 CXCL10** | TCATTGGTCACCTTTTAGTG |
| **FM1 CXCL2** | GGGTTGACTTCAAGAACATC | **FM1 IL-6** | GCAGAAAAAGGCAAAGAATC |
| **BM1 CXCL2** | CCTTGCCTTTGTTCAGTATC | **BM1 IL-6** | CTACATTTGCCGAAGAGC |
| **FM1 CXCL10** | AAAAAGGTCTAAAAGGGCTC | **FM1 TNF** | AGGCAGTCAGATCATCTTC |
| **BM1 CXCL10** | AATTAGGACTAGCCATCCAC | **BM1 TNF** | TTATCTCTCAGCTCCACG |
| | | **FM1 TGFb1** | AACCCACAACGAAATCTATG |
| | | **BM1 TGFb1** | CTTTTAACTTGAGCCTCAGC |
| | | **FM1 XCL1** | TACATTGTGGAAGGTGTAGG |
| | | **BM1 XCL1** | TGGTGTAGGTCTTGATTCTG |
| | | **FM1 YWHAZ** | AACCTTGACATTGTGGACATC |
| | | **BM1 YWHAZ** | AAAACTATTTGTGGGACAGC |

#### Supplementary Figure Legends

**Figure S1:**

IncuCyte assay analysis of total cell number and % cell death measured using Sytox Green uptake for indicated ovarian cancer cells. Cells were treated with; Control (vehicle), CHK1i (1 μM SRA737), LDHU (low-dose HU, 0.2 mM), HDHU (high-dose HU, 2 mM) or the indicated combinations. The data are the mean and SD of 6-30 determinations. These are representative of 2-3 experiments.

**Figure S2:**

IncuCyte assay analysis of total cell number and % cell death measured using Sytox Green uptake for indicated ovarian cancer cells. Cells were treated with: control (vehicle) or CHK1i (1 μM SRA737) + LDHU (low-dose HU, 0.2 mM. The data are the mean and SD of 6-30 determinations. These are representative of 2-3 experiments. These data were used to produce the % viable cell data in Figure 1C.

**Figure S3:**

CHK1i combination induces DNA damage and replication stress in ovarian cancer cells. Ovarian cancer cell lines and patient derived cells were treated with or without the CHK1i combination (1 μM SRA737 + 0.2 mM HU) for 24 hours, followed by cell harvesting and Western blot analysis of the indicated markers. The band intensities relative to the control for each cell line are shown for each band. The upper and lower numbers are for the respective treated bands for RPA2. This is representative of three individual experiments.

**Figure S4:**

Depletion of $CD8^+$ T cell. Representative plots of FACS analysis to confirm $CD8^+$ T cell depletion in blood at the CHK1i+LDHU treatment endpoint. Bars represent the mean ± SD. Statistical analysis was performed by One-way ANOVA. The level of $CD8^+$ T cells in the peripheral blood of mice harvested at the endpoint of the experiment shown in Figure 6D.

Supplementary Figure S1

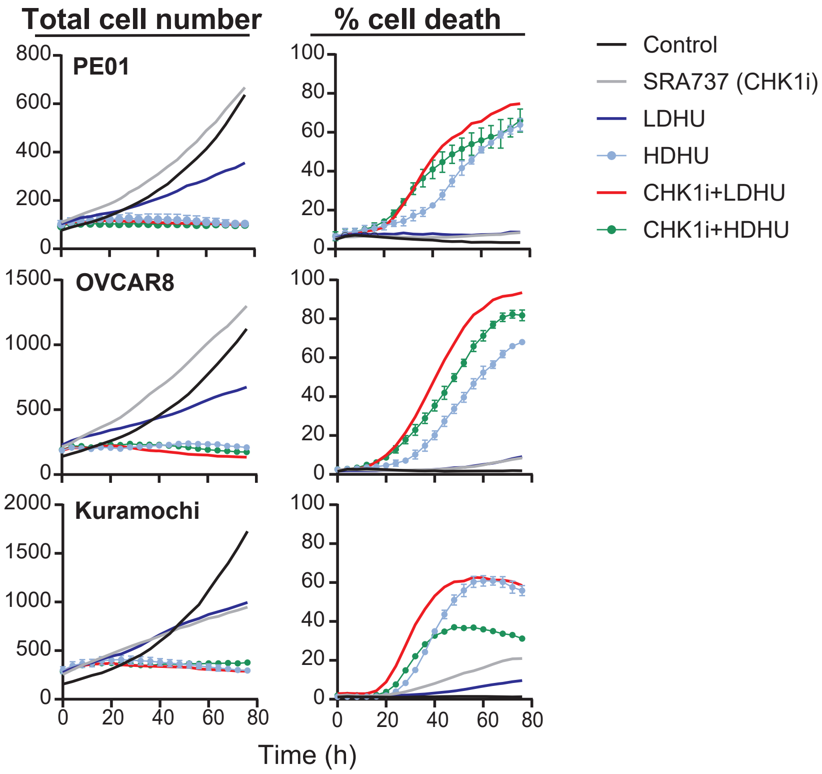

Supplementary Figure S2

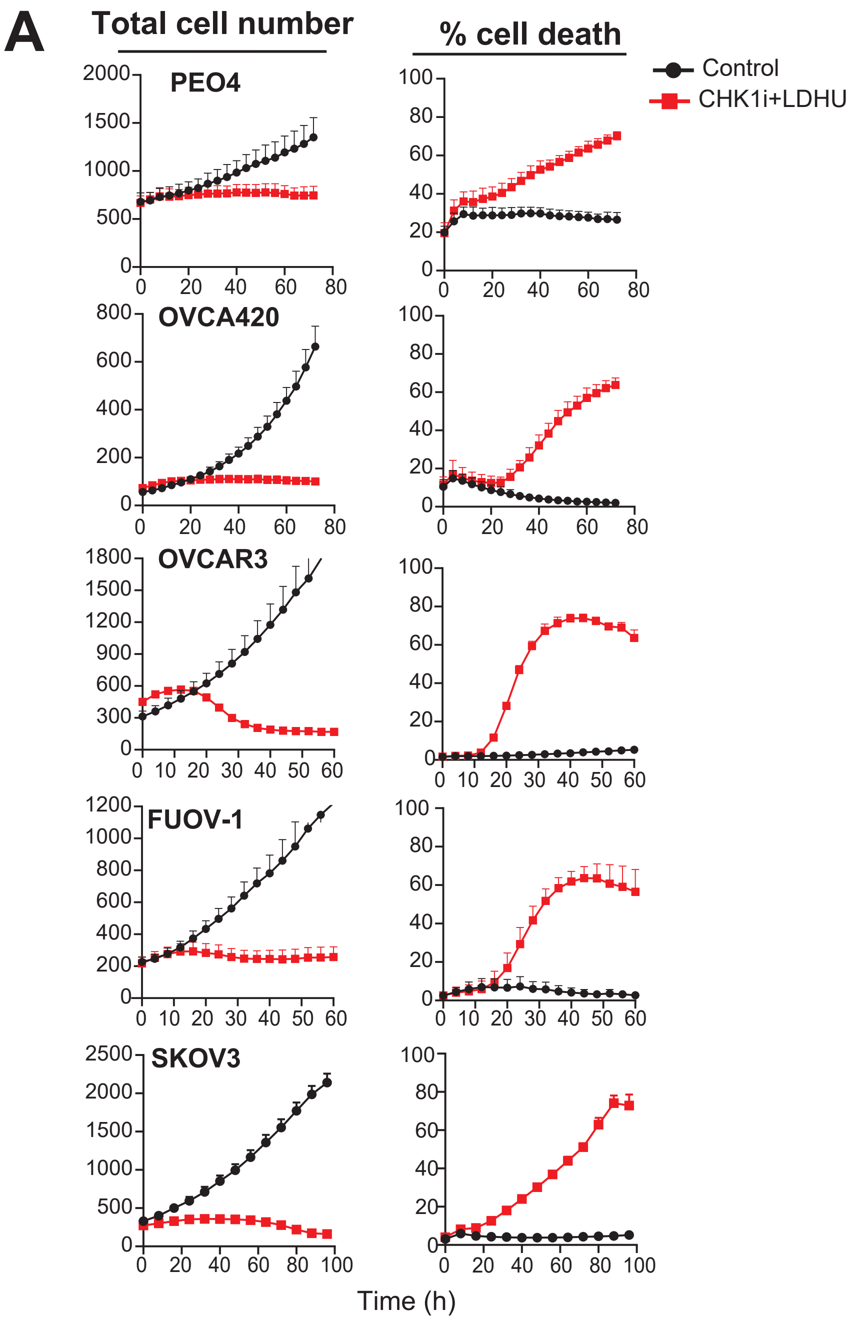


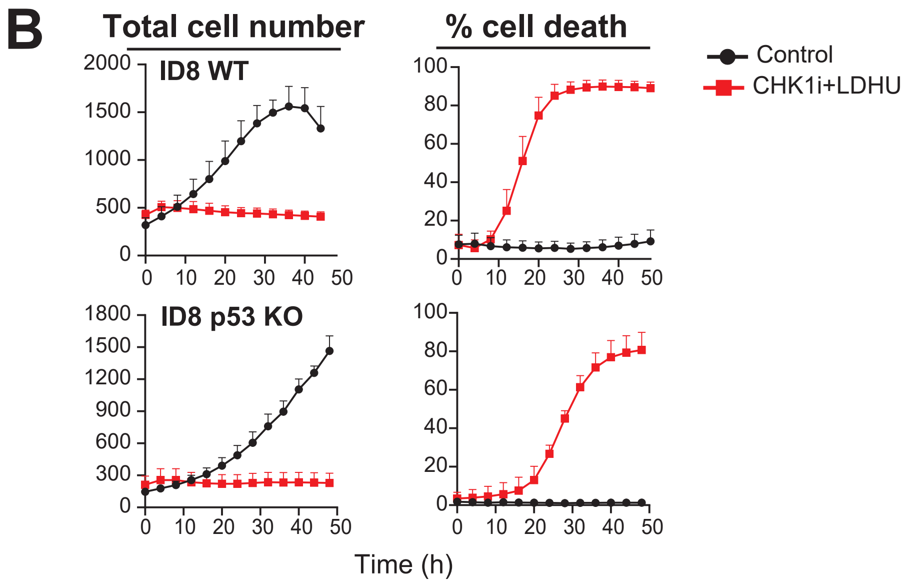

Supplementary Figure S3

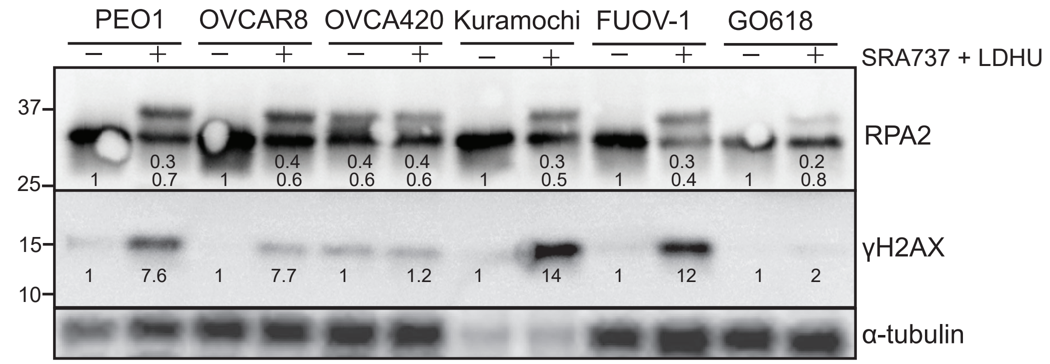

Supplementary Figure S4

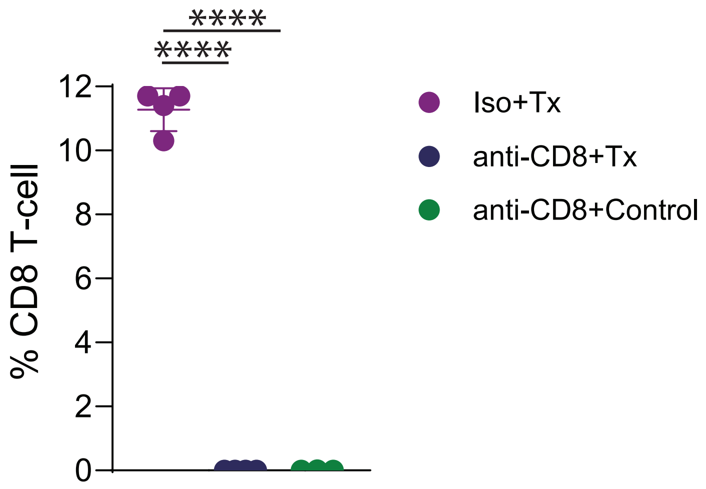
